# Myofilament Glycation in Diabetes Reduces Contractility by Inhibiting Tropomyosin Movement, is Rescued by cMyBPC Domains

**DOI:** 10.1101/2021.06.09.447778

**Authors:** Maria Papadaki, Theerachat Kampaengsri, Samantha K. Barrick, Stuart G. Campbell, Dirk von Lewinski, Peter P. Rainer, Samantha P. Harris, Michael J. Greenberg, Jonathan A. Kirk

**Author notes:** Corresponding Author Jonathan A. Kirk, Ph.D., Department of Cell and Molecular Physiology, Loyola University Chicago Stritch School of Medicine, Center for Translational Research, Room 522, 2160 S. First Ave., Maywood, IL 60153.

## Abstract

Diabetes doubles the risk of developing heart failure (HF). As the prevalence of diabetes grows, so will HF unless the mechanisms connecting these diseases can be identified. Methylglyoxal (MG) is a glycolysis by-product that forms irreversible modifications on lysine and arginine, called glycation. We previously found that myofilament MG glycation causes sarcomere contractile dysfunction and is increased in patients with diabetes and HF. The aim of this study was to discover the molecular mechanisms by which MG glycation of myofilament proteins cause sarcomere dysfunction and to identify therapeutic avenues to compensate. In humans with type 2 diabetes without HF, we found increased glycation of sarcomeric actin compared to non-diabetics and it correlated with decreased calcium sensitivity. Depressed calcium sensitivity is pathogenic for HF, therefore myofilament glycation represents a promising therapeutic target to inhibit the development of HF in diabetics. To identify possible therapeutic targets, we further defined the molecular actions of myofilament glycation. Skinned myocytes exposed to 100 μM MG exhibited decreased calcium sensitivity, maximal calcium-activated force, and crossbridge kinetics. Replicating MG’s functional affects using a computer simulation of sarcomere function predicted simultaneous decreases in tropomyosin’s blocked-to-closed rate transition and crossbridge duty cycle were consistent with all experimental findings. Stopped-flow experiments and ATPase activity confirmed MG decreased the blocked-to-closed transition rate. Currently, no therapeutics target tropomyosin, so as proof-of-principal, we used a n-terminal peptide of myosin-binding protein C, previously shown to alter tropomyosin’s position on actin. C0C2 completely rescued MG-induced calcium desensitization, suggesting a possible treatment for diabetic HF.

## Introduction

Almost half a billion people worldwide have type 2 diabetes mellitus (T2DM), a figure that is expected to increase by ~40% within 25 years [1]. Diabetes doubles the risk of developing heart failure [2, 3], independent of the effect on microvascular disease and hypertension. Left unchecked, the rapid expansion of diabetes will result in an explosion of heart failure cases. Therefore, it is critical that we discover the underlying molecular mechanisms that link these two conditions so they might be targeted therapeutically. We previously reported that one such link is likely mediated by methylglyoxal (MG) [4], a reactive carbonyl species that is formed from the degradation of triose phosphates [5] during glycolysis. MG can rapidly and irreversibly react with arginine and lysine amino acids on proteins [6], a process called glycation. Glycated proteins are known as advanced glycation end-products (AGE), and can bind to Receptors of AGE on the cell surface [7], inducing a signalling cascade resulting in oxidative damage and inflammation [8]. However, it is also possible for glycation of these residues to act as post-translational modifications that directly alter protein function, which is the mechanism explored in this study.

In recent years we and others have provided evidence that methylglyoxal may play a role in the development of heart failure [7, 9] through glycation of intracellular proteins involved in excitation-contraction coupling. For example, methylglyoxal can react with Ryanodine Receptor and SERCA in the hearts of type I diabetic rats and alter intracellular calcium handling [10, 11]. Our previous work, however, was the first to make the connection in humans, showing that MG modifications were increased in the cardiac myofilament of patients who had diabetes and heart failure, but not in heart failure patients without diabetes or healthy patients [4]. These MG modifications, occurring primarily on actin and myosin, reduced cardiomyocyte myofilament calcium sensitivity and maximal calcium-activated force [4]. However, whether MG modifications precede the development of overt systolic dysfunction, and thus represent a possible cause of increased heart failure risk in diabetic patients, is unknown. Genetic mutations in sarcomere proteins that reduce calcium sensitivity are a cause of cardiomyopathy and heart failure [12]. Thus, if myofilament glycation is elevated by diabetes and similarly reduce calcium sensitivity, they are likely pathogenic for the development of heart failure.

Since methylglyoxal modifications are irreversible, there are two possible approaches to correct the dysfunction: 1) decrease MG levels so these harmful adducts are not formed in the first place, or 2) identify treatments that can compensate for the dysfunction despite the continued presence of the MG modifications. Genetic approaches to depress MG levels in animal models may be beneficial, for example a recent study showing that AAV overexpression of Glo1, the enzyme that catalyzes MG, in endothelial cells improves function in a rat model of type 1 diabetes [13]. Unfortunately, approaches to reduce MG levels clinically have already failed. Specifically, compounds called “AGE-breakers”, drugs aimed at reducing levels of MG and AGE [14, 15] showed no benefit in clinical trials. In fact, one trial in patients with type 2 diabetes was terminated early due to a negative impact [16]. Our therapeutic strategy aims at correcting the dysfunction occurring at the myofilament level, since the myofilament is a highly tuneable system [17] that is likely amenable to this approach.

In this study, we show in diabetic human hearts without heart failure that MG modifications are already increased compared to non-diabetics and correlate with early myofilament dysfunction. A successful strategy for restoring function would need to be informed by first discovering the molecular mechanism(s) by which MG inhibited myofilament function. Therefore, we measured the impact of MG on myofilament kinetics, used a computer model of myofilament activation to predict molecular changes, and validated the predictions through biophysical assays. These comprehensive approaches revealed that MG decreases the rate constant for the transition of tropomyosin from its blocked state to its closed state on actin, effectively making it harder to activate the thin filament. The n-terminal domain of cardiac myosin binding protein C (cMyBPC) has been shown to activate the thin filament by altering tropomyosin’s position on actin [18]. Indeed, the effects of MG on myofilament calcium sensitivity were rescued by recombinant cMyBPC domains, suggesting that targeting thin filament activation could be a viable therapeutic strategy to break the connection between diabetes and heart failure.

## Methods

Expanded Methods are presented in Supplemental Material

### Human and animal studies

Human left ventricular tissue samples were obtained at the Medical University of Graz from organ donors whose hearts were rejected for transplantation but did not have heart failure. Human study protocols were approved by the Ethics Committee at the Medical University of Graz (28-508 ex 15/16). All patients gave informed consent and the investigation conformed to the principles outlined in the declaration of Helsinki. Animal studies were approved by the Loyola University Chicago Institutional Animal Care and Use Committee (IACUC number 2019029) according to the NIH Guide for the Care and Use of Laboratory Animals. C57/Bl6J male mice 2-4 months of age were purchased from Jackson laboratories (Jackson labs, USA). Animals were euthanised by placing them in an induction chamber with isoflurane vaporizer (5%). The animal remained in the chamber until unconscious, as determined by corneal reflex. Following isofluorane exposure, animals were euthanised by cervical dislocation and heart extubating.

### Mass spectrometry

Myofilament samples were prepared as stated previously [4]. Samples were run on a 4-12% SDS- PAGE gel, stained with Coomassie stain, actin and myosin bands excised, and then de-stained overnight. Gel bands were prepared for mass spectrometry by digesting them in 2 μg/band Trypsin/LysC protease mix (Thermo Scientific) for 16 hours at 37 °C. 200 - 300 ng of peptides were loaded onto an UltiMate 3000 nanoHPLC coupled to a LTQ Orbitrap XL (Thermo). MS data analysis was performed using the Peaks Bioinformatics Software. For analysis, glycated peptides were normalized to the corresponding total peptides for each sample.

### Functional assessments and recombinant proteins

Skinned myocytes were prepared as previously described [4, 19]. ATPase activity and tension cost were measured in skinned fibers using an in-house system, as previously [20, 21]. Cardiac myosin S1 and actin and tropomyosin were expressed and purified as described in the Supplemental methods. Tropomyosin blocked-to-closed state transition (KB) and ADP release were measured using stopped-flow methods [22] (see Supplementary methods for details).

### Computational modelling

A previously published model of myofilament Ca^2+^ activation [24] was used to identify molecular changes to sarcomeric proteins that could plausibly explain observed effects of methylglyoxal on skinned cardiac fibres. Baseline parameters from the published model were used as a starting parameter set. The model was implemented and run in MATLAB (Mathworks) as previously described.

### Statistical analysis

Data are presented as mean ± standard error. Quantitative data were analysed using Prism 8 (GraphPad software) and stopped-flow data were analysed using MATLAB. All statistical analyses were performed using Student’s t-test, paired Students t-test, chi-squared test, or two-way repeated measures ANOVA with Sidak post hoc test, depending on the data set, as indicated in the text. P<0.05 was considered significant.

## Results

### Diabetics have increased glycation that correlates with myofilament function

We previously showed that patients with diabetes and heart failure exhibited increased MG glycation of myofilament proteins, primarily actin and myosin [4]. Here, we first sought to measure these MG modifications in diabetics without heart failure, to test whether they might precede heart failure and represent a possible therapeutic target. We utilized left ventricular (LV) tissue from donor hearts without heart failure, and either with type II diabetes or without (Non-Diabetic, *n* = 9; Diabetic, *n* = 6, demographic and basic clinical data shown in **Table 1**).

**Table 1.** Human Subject Characteristics

|  | Non-Diabetic | Diabetic | P-value |
| --- | --- | --- | --- |
| n | 9 | 6 |  |
| Age (years) | 63 ± 7 | 61 ± 10 | 0.82 |
| % Female | 33 | 50 | 0.62 |
| BMI | 30 ± 7 | 29 ± 2 | 0.79 |
| % Diabetes | 0 | 100 | 0.0001 |
| % Ischemia | 0 | 0 | N/A |
| LVEF (%) | 66 ± 7 | 66 ± 4 | 0.92 |
| % Hypertension | 22 | 100 | 0.007 |
| Anti-hypertensive drugs (%) | 16 | 100 | 0.001 |
| Oral anti-diabetics (%) | 0 | 66 | 0.01 |
| $\beta$ blockers (%) | 16 | 50 | 0.23 |

BMI, body mass index; LVEF, left ventricular ejection fraction. Statistics by Student's t-test or Chi-squared test.

For each of these LV samples, the myofilament was enriched, separated by gel electrophoresis, and the actin and myosin bands excised and analysed for MG glycation by mass spectrometry. Diabetic patients exhibited significantly elevated glycation (including glyoxal and methylglyoxal-derived modifications) on sarcomeric actin compared to Non-Diabetic hearts (**Figure 1A**, **Supplemental Table 1**). However, no differences were observed in myosin glycation between the groups (**Supplemental Figure 1A, Supplemental Table 2**).

**Figure 1:**
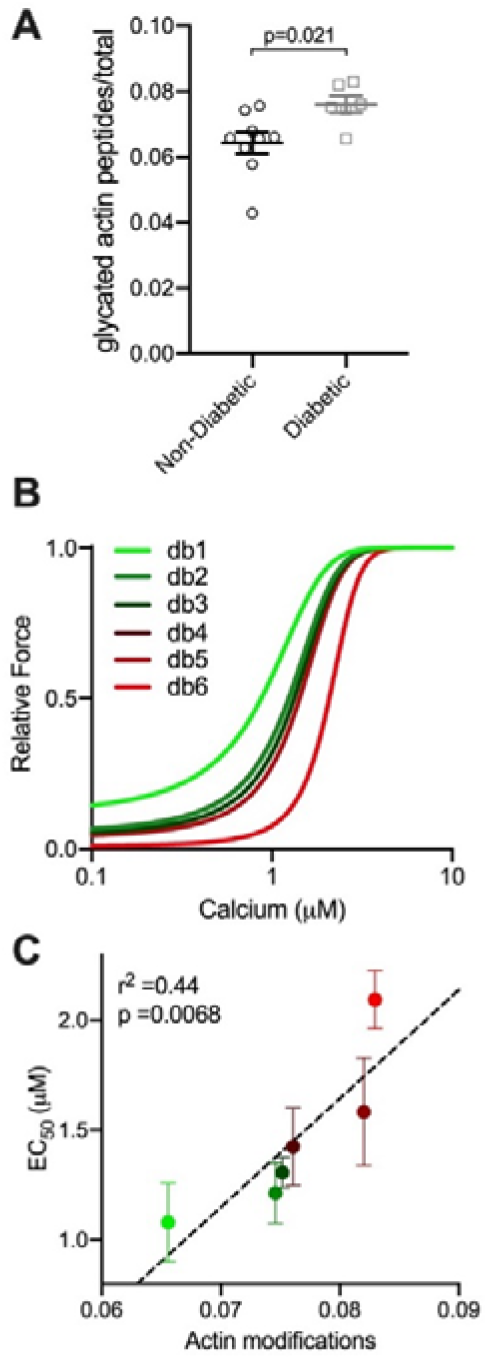
Glycation of sarcomeric actin was increased in Diabetic compared to Non-Diabetic subjects and correlates with decreased myofilament calcium sensitivity. (**A**) Glycated peptides normalized to total peptides as measured by mass spectrometry on actin, from Non-Diabetic (black circles, *n*=9) and Diabetic (grey squares, *n*=6) left ventricular tissue. Statistical comparison by Student’s t-test. (**B**) Mean fitted curves for force as a function of calcium concentration in human skinned myocytes isolated from Diabetic subjects (n=3-4 myocytes per subject, n=6 subjects) for the subject with the lowest (db1, green) to highest level of actin glycation (db6, red), showing progressive desensitization to calcium in subjects with increased glycation. (**C**) Mean EC_50_ as a function of total actin glycation detected by mass spectrometry for each human Diabetic sample (circles colour coded from the lowest glycation level, bright green to highest glycation level, bright red, *n*=6). The data are described by a positive correlation (dashed line, r^2^=0.44, p= 0.0068 by linear regression.

We next measured force-calcium relationships in skinned myocytes isolated from LV tissue from each Diabetic heart (n = 3 – 4 myocytes per heart) to understand the impact of increased actin glycation on sarcomere function. For each Diabetic heart, the average fitted curve for all myocytes measured from that heart are shown in **Figure 1B**, from the heart exhibiting the lowest level of actin glycation (db1, green line) to the highest level (db6, red line). A progressive rightward shift in the force-calcium relationship (calcium desensitization) was observed with increasing levels of actin glycation. This correlation was confirmed when mean calcium sensitivity of all cardiomyocytes from each subject was plotted against the level of actin glycation for each subject (**Figure 1C**). The strong positive correlation (r^2^ = 0.44, p = 0.0068 for a non-zero slope by linear regression), indicates that in diabetic hearts, increased levels of actin glycation are associated with a progressive decrease in myofilament calcium sensitivity (increased EC_50_).

Not all diabetic patients eventually develop heart failure, however. Indeed, while we observed increased actin glycation overall, some subjects had levels equal to Non-Diabetics. As such, some subjects exhibited normal myofilament function, so that no overall differences were observed in maximal calcium-activated force (F_max_) or calcium sensitivity (EC_50_) between Non-Diabetic versus Diabetic samples (**Supplemental Figure 1B-D**). Myofilament calcium desensitization does not always result in concurrent observable global dysfunction [23], but is known to be pathogenic for dilated cardiomyopathy and heart failure. Thus, in diabetics with increased glycation, MG modifications represent a possible therapeutic target to reduce the risk of developing heart failure in the progression of diabetic cardiomyopathy.

### Computational predications of molecular processes dysregulated by MG

Our next goal was to determine the molecular mechanism(s) of action of MG, as myofilament functional parameters like calcium sensitivity are an aggregate measurement of numerous molecular processes [24]. To predict which of these processes are modified by MG we used a multiscale computer model of cooperative myofilament activation [25]. To adequately constrain the model, it is necessary to know the functional impact of MG on both steady-state and kinetic parameters.

To measure these parameters, we used skinned myocytes isolated from LV tissue from three-month-old male C57Bl6/j mice treated with 100 μM MG for 20 mins. In healthy tissue, levels of free MG are around 1 – 10 μM [6, 26, 27] and approximately double in disease [28]. However, the majority of MG is found on glycated proteins [29], meaning these measurements of free MG significantly under-estimate the amount of MG glycation that occurs. To ensure this MG treatment increased glycation of the myofilament, we treated mouse left ventricular skinned myocytes with 100 μM MG for 20 mins. We then solubilized these samples, ran them on SDS PAGE gels, excised actin and myosin bands, and analysed them via high resolution nHPLC-MS/MS. We found that MG modifications on actin and myosin were increased after the *in vitro* MG treatment (**Supplemental Figure 2 and Supplementary Table 3, 4**).

The impact of MG on steady-state parameters is described by the force-calcium relationship. As we had shown previously [4], exposure to MG decreased both calcium sensitivity and F_max_ (**Figure 2A-C**). To determine the effect on kinetics, we measured the rate of force recovery after a slack-restretch maneuver (*k*_tr_) before and after MG treatment. Further, *k*_tr_ was measured when the myocyte was exposed to either maximal (46.8 μΜ) or submaximal (1.7058 μΜ, approximately EC_50_) calcium. Treatment with 100 μM MG decreased *k*_tr_ at maximal and sub-maximum Ca^2+^ activation (**Figure 2D-F**).

**Figure 2:**
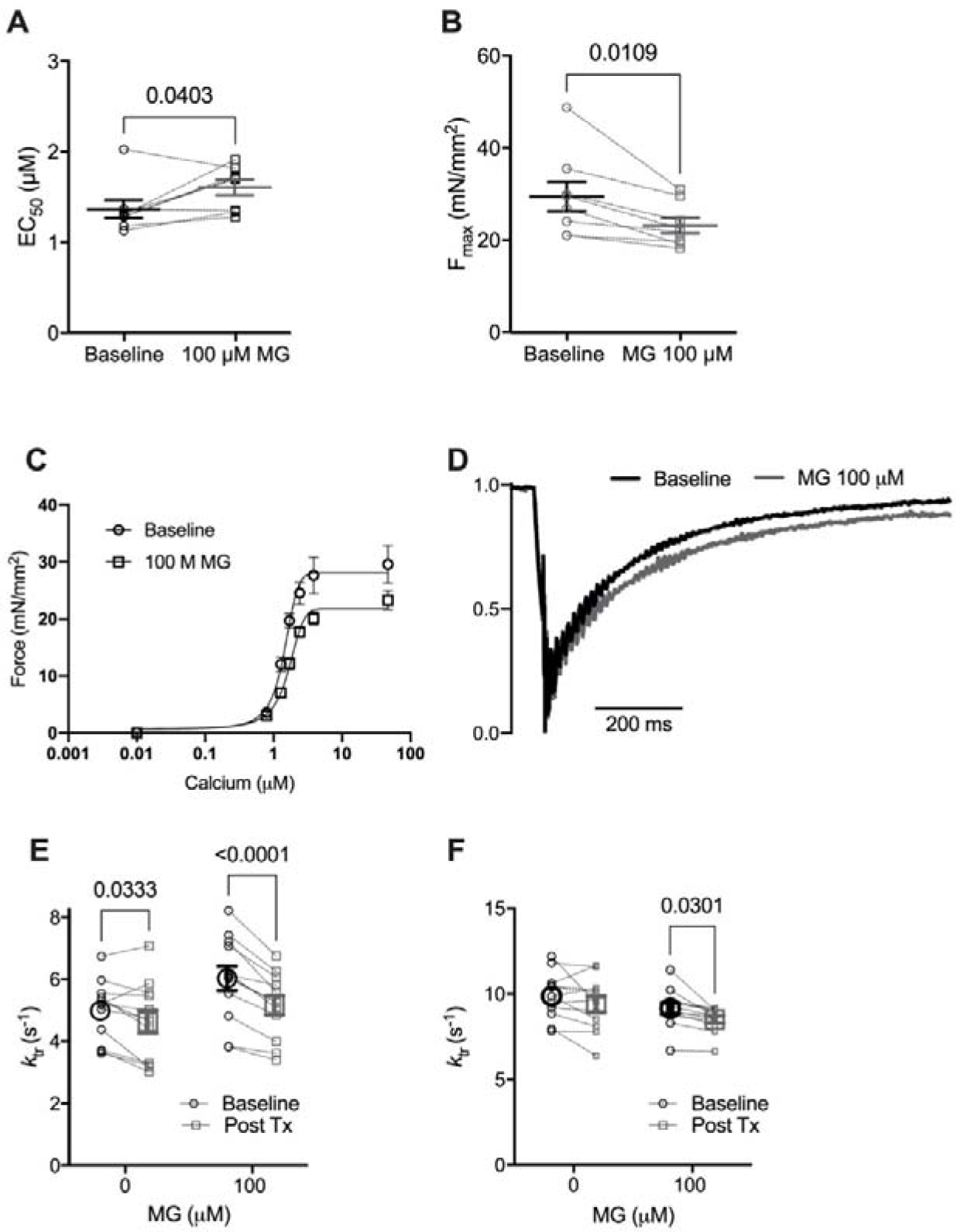
Methylglyoxal decreases calcium sensitivity, maximal calcium-activated force and rate of force redevelopment (*k*_tr_). (**A**) Individual and mean ± SEM calculated EC_50_ (panel **A**) and F_max_ (panel **B**). (**C**) Mean force as a function of calcium concentration and fitted curves for mouse skinned myocytes before (circles, solid line) and after (squares, dashed line) exposure to 100 μM Methylglyoxal (MG). (**D**) Normalized *k_tr_* curve before (black) and after (grey) treatment with 100 μM MG at submaximal Ca^2+^ concentration (average from 12 cells). (**E**) Individual and mean ± SEM calculated *k*_tr_ values before (Baseline, black circles) or after (Post Tx, grey squares) treatment with 0 (no treatment) or 100 μM MG at sub-maximal Ca^2+^ concentration (*n*=12 myocytes from 4 mice, p_interaction_ = 0.016) (Mean ± SEM). (**F**) *k*_tr_ before (black circles) or after (grey squares) treatment with 0 or 100 μM MG at maximal Ca^2+^ concentration (*n*=12 myocytes from 4 mice, p_interaction_ = n.s.). Statistical comparisons made with 2-way repeated measures ANOVA with Sidak post-hoc test.

Next, using the computational model of myofilament activation, individual model parameters were adjusted in an attempt to recapitulate the differences between the baseline and MG-treated functional data (**Figure 3A**). The model was first used to generate baseline force-calcium (blue line, **Figure 3B**) and *k*_tr_-calcium (blue line, **Figure 3C**) relationships, using previously published baseline model parameters [25]. We then altered individual or combinations of parameters to recapitulate the simultaneous impact on EC_50_ (**Figure 2A**), F_max_ (**Figure 2B**) and *k*_tr_ (**Figure 2D-F**).

**Figure 3:**
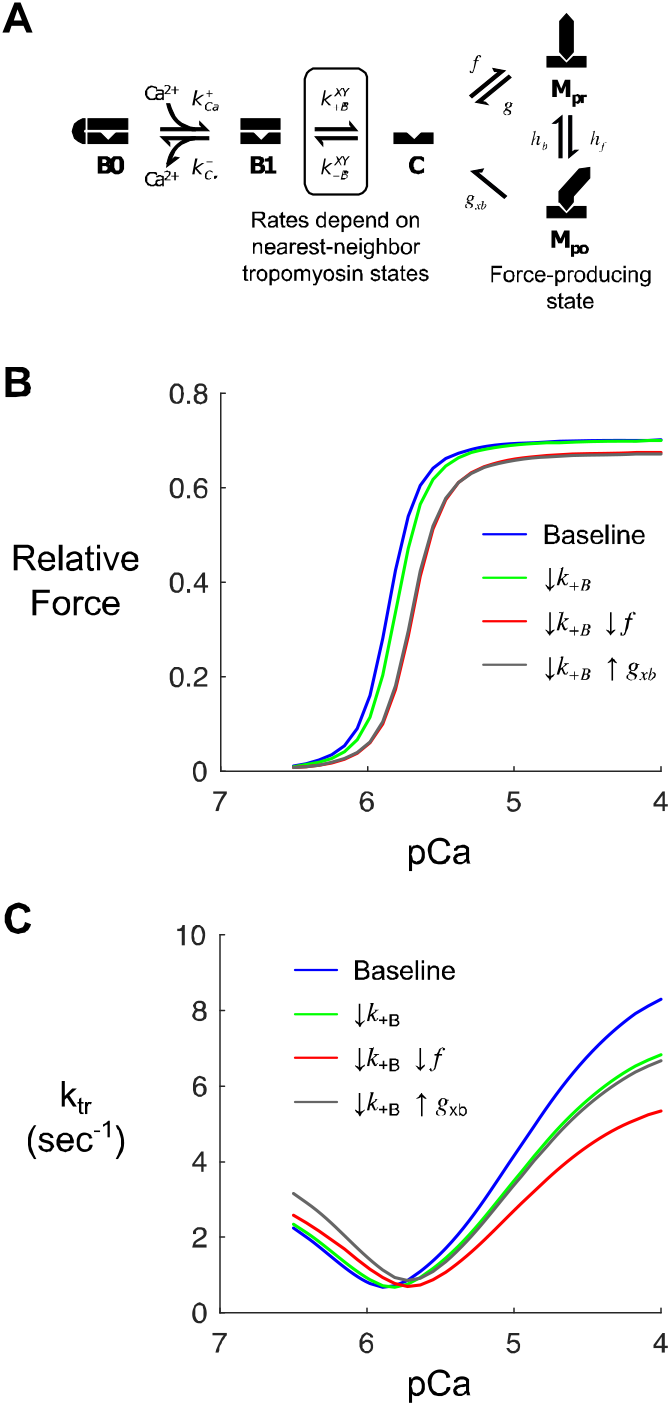
Computer modelling to identify potential methylglyoxal molecular effects. (**A**) Kinetic scheme for a previously published model of myofilament Ca^2+^ activation [24]. The model depicts states of thin filament regulatory units (B0, blocked and Ca^2+^-free; B1, blocked and Ca^2+^-bound; C, closed) and cycling of associated myosin crossbridges (M_pr_, myosin pre-powerstroke; M_po_, myosin post-powerstroke). (**B**) Steady-state force-pCa curves produced by the model under baseline (control) conditions and various perturbations intended to qualitatively mimic observed effects of MG. Simulations show the anticipated effects of slowing the blocked-closed transition of tropomyosin (*k_+B_*) on its own (green) or in combination with slowing myosin attachment (*f*, red) or myosin detachment (*g*, gray). (**C**) Model-predicted k_tr_-pCa relationships corresponding to the same conditions as in panel B.

The model indicated that simultaneous changes in two parameters were sufficient to reproduce this behaviour: 1) a ~10% decrease in the tropomyosin blocked-to-closed rate transition (*K*_B_) and 2) a decrease in the myosin crossbridge duty cycle from either a 20% decrease in the myosin binding rate (*f*) or a 30% increase in the myosin detachment rate (*g_xb_*). The individual and combined effects of these changes are shown in green, red, and grey on **Figure 3 B, C**. We subsequently searched for experimental evidence that MG treatment affected these molecular processes.

### MG increases the probability of thin filaments being in the “blocked” state

Tropomyosin is found in three positions on actin: blocked, closed and open. These positions are determined by calcium and myosin binding to the thin filament [30–32]. The equilibrium constant describing the transition from the blocked to closed tropomyosin state (*K*_B_) was determined using stop flow experiments, where the rate of myosin binding to pyrene-labelled regulated thin filaments was measured at high (pCa 4) and low (pCa 9) calcium concentrations [22, 32]. The observed rates for myosin S1 binding to the thin filament at high and low calcium were used to calculate the equilibrium constant, *K*_B_ (see Supporting Materials for details). Since these experiments have not been previously performed, we needed to examine the effects on both the lower and the higher dose of MG on thin filaments. Stopped-flow experiments were performed using reconstituted regulated thin filaments before and after exposure to 10 μM MG for 20 mins. MG decreased *K*_B_ by approximately 50% in regulated thin filaments (**Figure 4A, B**). A decreased equilibrium constant means that the blocked state will be more favoured over the closed state. Although we cannot exclude the possibility that MG decreases K_B_ by increasing the rate of the closed-to-blocked transition, the decrease in K_B_ is consistent with the reduced rate of the blocked-to-closed transition predicted by the computational modelling.

**Figure 4:**
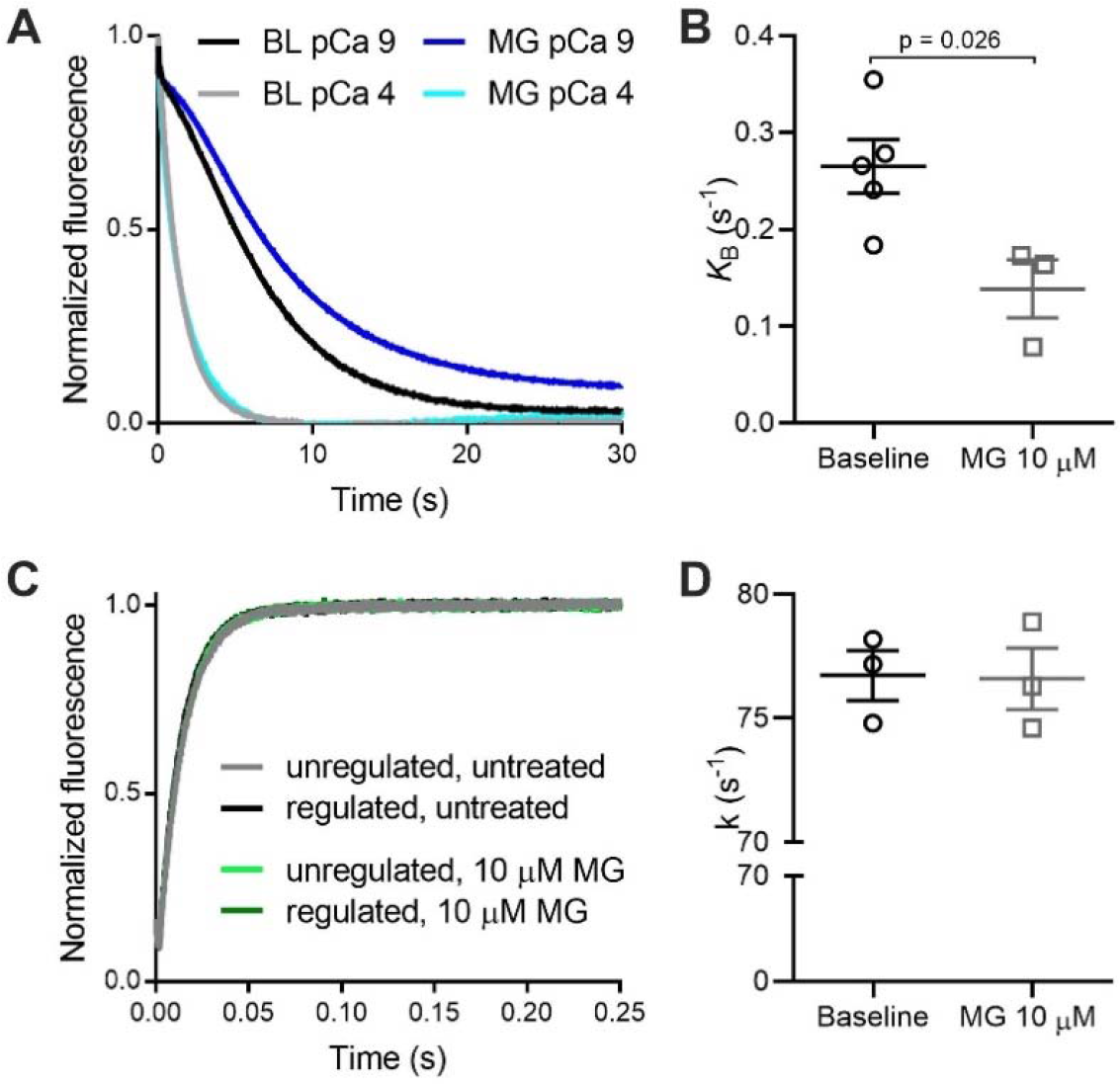
Methylglyoxal decreases the tropomyosin blocked-to-closed rate transition. (**A**) Normalized fluorescence of pyrene-labelled actin as myosin S1 binds to regulated thin filaments at pCa 4 (high calcium) and pCa 9 (low calcium). The pyrene fluorescence decreases upon S1 binding. Rate of myosin S1 binding to regulated thin filaments is higher at pCa 4 due to thin filament activation by calcium. The difference in the rates at pCa 4 and pCa 9 is used to calculate *K*_B_. The black and grey lines represent untreated filaments (BL) and the dark/light blue lines represent thin filaments treated with 10 μM MG. (**B**) Individual and mean ± SEM *K*_B_ values for untreated (baseline, black dots, n=5) or treated thin filaments with 10 μM MG (grey squares, n=3). (**C**) Measurement of the actomyosin detachment rate. Normalized fluorescence of pyrene-labelled actin as myosin S1 bound to ADP detaches from regulated thin filaments upon addition of ATP. The rate of actomyosin detachment is measured from the increase in pyrene fluorescence. Unregulated filaments (black, grey) or regulated filaments (dark/light green) treated or not with 10 μM MG. (**D**) Individual and mean ± SEM rate constant (*k*) for ADP release from myosin S1 binding to regulated thin filaments untreated (baseline, black circles) or treated with 10 μM MG (grey squares) (*n*=3 for each). Statistical comparisons by Students t-test.

The model also predicted MG either increases the crossbridge detachment rate or decreases the attachment rate. By stopped-flow, we can measure the detachment rate constant (k) of ADP release from actomyosin solution, which is the transition that limits crossbridge detachment rate at physiological ATP concentrations. We performed stopped-flow experiments by rapidly mixing 1) pyrene-labelled thin filaments pre-incubated with myosin S1 and ADP with 2) solution containing saturating ATP [33]. When thin filaments were incubated with 10 μM MG for 20 mins there was no difference in ADP release (**Figure 4C, D**). Even at the higher 100 μM dose, MG had no effect on the crossbridge detachment rate measured using reconstituted proteins in solution (**Supplemental Figure 3**).

To confirm the biochemical stopped-flow results in a more physiological system, we measured ATPase activity and active force production simultaneously in skinned mouse papillary fibers. Tension cost (the ratio of myosin ATPase activity to force production) is directly proportional to the number of post-power stroke crossbridges and therefore represents a measurement of the crossbridge detachment rate. We found that 40 mins incubation with MG decreased calcium sensitivity and F_max_, (as in skinned myocytes) as well as ATPase activity (**Figure 5**). The longer incubation time (40 minutes vs 20 minutes in skinned cells) was necessary because of the larger size of the fiber bundles. Since MG decreased both force and ATPase activity in parallel, there was no effect on tension cost (the ATPase activity to Force ratio), indicating that MG has no effect on the crossbridge detachment rate. By two measurements we confirm that MG has no effect on the crossbridge detachment rate, suggesting MG decreases the crossbridge attachment rate.

**Figure 5:**
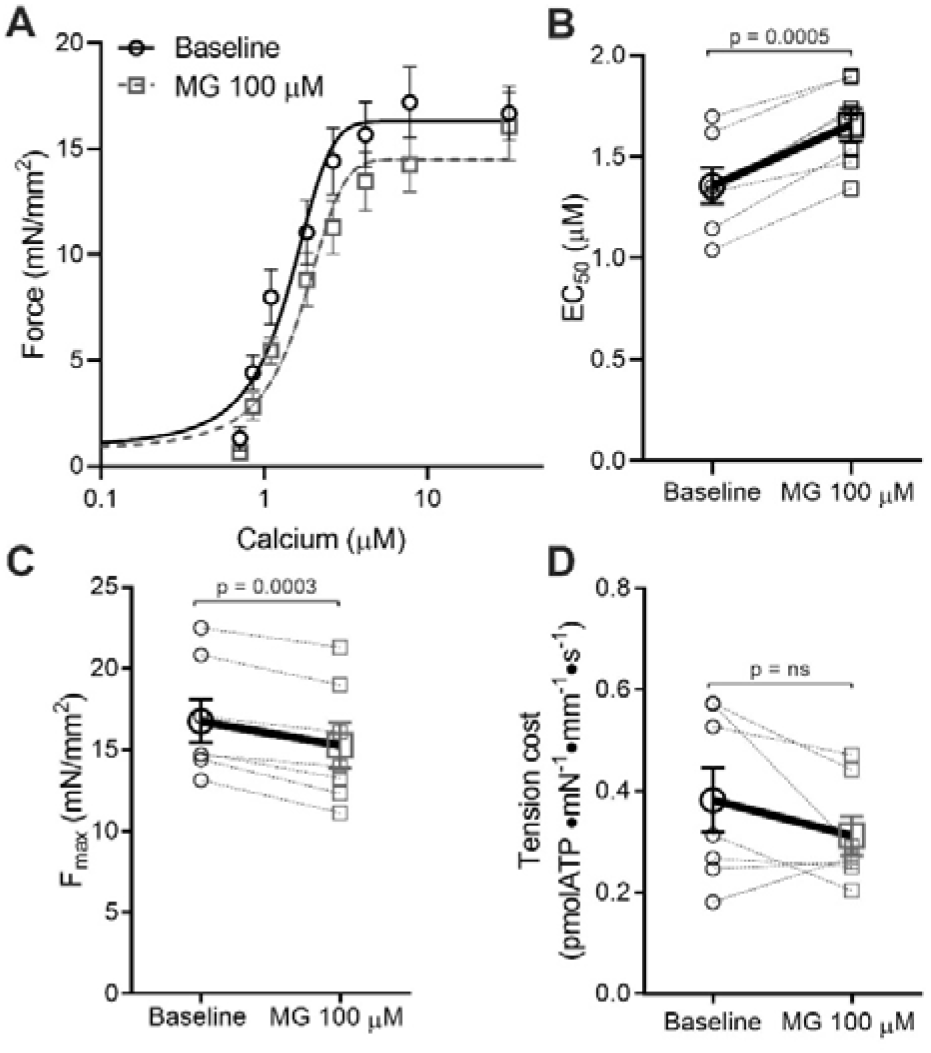
Methylglyoxal does not alter tension cost of mouse skinned fibers. (**A**) Average force-calcium data and fitted curves for skinned fibers before (black circles, solid line) or after (grey squares, dashed line) treatment with 100 μM MG for 40 mins (*n*=7 fibers from 4 mice). (**B**) Individual and mean ± SEM calculated EC_50_ values before (baseline, black circles) or after 100 μM MG (grey squares). (**C**) F_max_ before (baseline, black circles) or after 100 μM MG (grey squares) (*n*=7 fibers from 4 different mice). (**D**) Tension cost was calculated via the ratio of ATPase activity over force, before (baseline, black circles) or after 100 μM MG (grey squares). Statistical comparisons made by student’s paired t-test.

### N-terminal MyBPC fragment rescues MG-induced dysfunction but OM does not

Having established that MG depresses myofilament function by inhibiting tropomyosin movement on actin, we next aimed to determine whether we could rescue calcium sensitivity be specifically targeting these mechanisms. However, we first wanted to confirm that reducing MG itself is not an effective therapy. We used aminoguanidine (AG), an agent that has been reported to scavenge MG and prevent it from forming further modifications [34]. Skinned myocytes were pre-treated with 100 μM MG for 20 mins, a force-calcium relationship was measured, then treated with 1 mM AG for 5 mins and a second force-calcium relationship measured. This dose of AG and treatment time was chosen because previous experiments in intact isolated cardiomyocytes showed it was capable of improving function [35]. However, we found that AG had no effect on calcium sensitivity or F_max_ (**Supplemental Figure 4)**, indicating it was unable to reverse the deleterious functional effects of MG.

Next, we supposed that calcium sensitizers that did not target tropomyosin might be ineffective at rescuing MG-induced dysfunction. Thus, we first used Omecamtiv Mecarbil (OM), a myosin activator which increases Ca^2+^ sensitivity in isolated cardiomyocytes [36]. We hypothesized that MG’s impact on tropomyosin would block the OM-activated myosin heads from binding actin, and thus not be rescued with this treatment. We found that in the absence of MG, treatment with 1 μM OM for 2 mins increased calcium sensitivity and decreased F_max_ as previously [36]. However, in skinned myocytes pre-incubated with 100 μM MG for 20 mins, OM had no effect on Ca^2+^ sensitivity and further reduced F_max_ (**Figure 6A,B)**. These results indicate that OM is not effective in reversing the functional defects by MG, possibly even exacerbating the dysfunction by further reducing F_max_, supporting MG’s action through tropomyosin.

**Figure 6:**
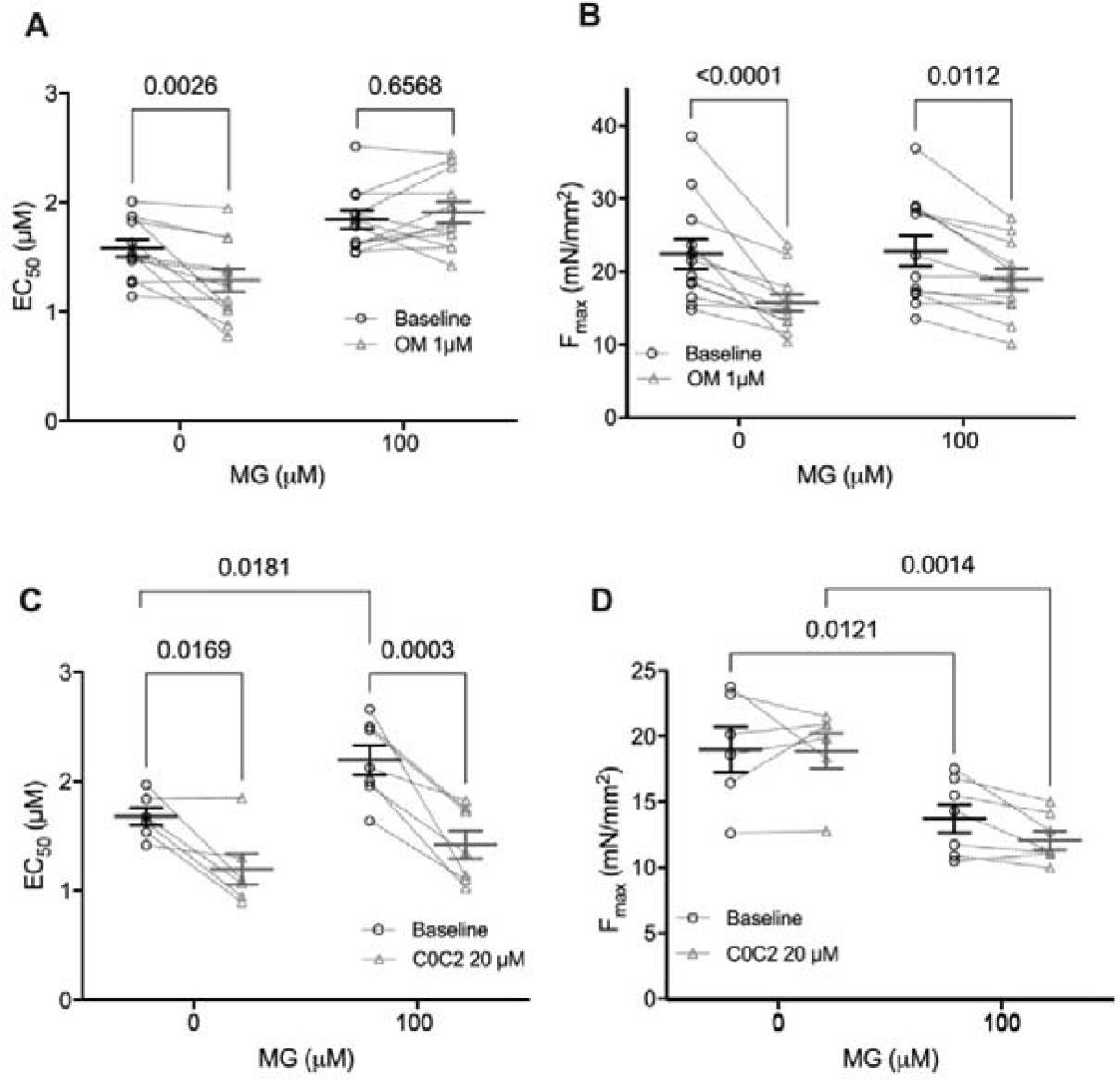
C0C2 rescues the calcium sensitivity decrease caused by MG in skinned cardiomyocytes but OM does not. (**A**) Individual and mean ± SEM EC_50_ values from skinned myocytes pre-exposed to 0 μM MG (*n*=6) or 100 μM MG (*n*=7 from 3-4 mice) before treatment with cardiac myosin binding protein C peptide C0C2 (baseline, black circles) or after (C0C2 20 μM, grey triangles). Paired before and after data for an individual cell are connected by dashed lines. (**B**) Individual and mean ± SEM F_max_ values from skinned myocytes pre-exposed with 0 μM MG (*n*=4) or 100 μM MG (*n*=7 from 3 mice) before with C0C2 (baseline, black circles) or after (C0C2 20 μM, grey triangles). Individual and mean ± SEM calculated EC_50_ (panel **G**) and F_max_ (panel **F**) from skinned myocytes pre-exposed to 0 μM MG (*n*=8) or 100 μM MG (n= 12 from 4 mice) before (baseline, black circles) or after treatment with 1 μM Omecamtiv Mecarbil (1 μΜ OM, grey triangles) (*n*=12 myocytes from 4 mice). Paired before and after treatment data for each individual cell are connected by dashed lines. Statistical comparisons by two-way repeated measures ANOVA with Sidak post-hoc test.

Knowing that MG increases the population of thin filaments in the inhibitory blocked state, we next aimed to discover whether a molecule that activates the thin filament could reverse the action of MG on the sarcomere. Currently, there are no small molecules that act on tropomyosin [37–39], so we used recombinant cMyBPC C0C2 peptide, which has been shown to activate the thin filament by pushing tropomyosin into positions that are more permissive for myosin binding to the thin filament [40]. The cMyBPC C0C2 peptide consists of the 466 n-terminal amino acids that encompass the cardiac specific C0 domain, two Ig domains (C1, C2), a proline-alanine linker between C1 and C2, and a unique cMyBPC “motif” or M-domain that connects C1 and C2. We hypothesized that C0C2 would be able to reverse the functional defects of MG through likewise displacing tropomyosin.

In mouse skinned myocytes not exposed to MG, incubation with 20 μM C0C2 for 20 minutes increased calcium sensitivity with no change in F_max_ (**Figure 6C, D**), which is in line with published data [41, 42]. At this concentration of C0C2, it is known to interact with actin [42, 43] and is thus likely to impact tropomyosin’s position on actin as intended. Skinned myocytes pre-incubated with 100 μM MG were significantly desensitized to calcium (p = 0.036 by two-way ANOVA) and trended towards a decrease in F_max_ (p = 0.052 by two-way ANOVA), recapitulating our prior data. When these MG-preincubated myocytes were treated with C0C2 for 20 minutes, Ca^2+^ sensitivity was restored (from EC_50_= 2.19 to 1.42, p=0.0018 by two-way ANOVA, *n* = 7, **Figure 6C**). While there was a significant decrease in F_max_ with C0C2 treatment in the MG-exposed myocytes (from F_max_ = 13.9 to 12.2, p=0.043 by two-way ANOVA, *n* = 7, **Figure 6D**), the magnitude of this decrease was less than 15% of the pre-treatment level and is likely due to normal myocyte rundown during multiple activations. Furthermore, there was no interaction between C0C2 and MG treatments on the effect of F_max_ (p = 0.57 by two-way ANOVA). Overall, these data show that C0C2 can reverse the depressed calcium sensitivity caused by MG, presumably by moving tropomyosin on actin into a position that is favourable for myosin contraction.

## Discussion

We have shown for the first time that diabetics free of heart failure exhibit increased MG actin glycation compared to those without diabetes. In diabetics, the magnitude of elevated glycation positively correlated with myofilament desensitization to calcium. Not all diabetics eventually develop heart failure, so it is reasonable that in this cohort we would detect a range of glycation and function. It is thus unlikely that overt dysfunction could be used to predict the eventual development of heart failure in diabetics, and MG modifications may represent a more sensitive biomarker. We hypothesize those subjects with high levels of glycation and calcium desensitization would be the most likely to develop diabetic cardiomyopathy and heart failure [2]. As these were donor hearts, it is impossible to know for sure whether they would have eventually developed cardiac dysfunction, however, point mutations that cause familial dilated cardiomyopathy frequently cause similar myofilament calcium desensitization that eventually leads to heart failure [12]. Thus, the cellular dysfunction induced by this glycation represents a possible therapeutic target for inhibiting the development of diabetic cardiomyopathy [44], which can lead to heart failure [45].

In non-failing diabetics, the median magnitude of the increase in actin glycation was about half of what we previously observed in diabetic patients with heart failure (~20% vs ~40% increase compared to controls [4]). This relationship suggests that myofilament MG glycation increases in parallel with disease severity, preceding global cardiac dysfunction. Importantly, while our previous work was in diabetic heart failure patients [4], these findings in otherwise healthy diabetics decouple the impact of myofilament glycation on cardiomyocyte function from adaptive and maladapting processes occurring during end-stage heart failure. That the increased glycation was detected only on actin, and not myosin, was unexpected but suggests a preference for MG glycation. While no previous studies have identified any site specificity for MG glycation, these data indicate there are certainly preferred targets, possibly due to length or concentration of MG exposure as well as the accessibility and reactivity of the modified residues. The increased levels of MG glycation on solely actin are still harmful as our previous work showed dysfunction was induced by MG glycation of actin or myosin independently [4].

Unfortunately, broadly targeting the MG-induced dysfunction with compounds that inhibit MG modifications or general systolic activators was unsuccessful in reversing the dysfunction. A combination of biophysical functional approaches, computer simulation, and stopped-flow experiments revealed that MG modifications increase the population of the tropomyosin blocked state, therefore inhibiting myosin from binding to the thin filament and initiating contraction. Effectively, MG makes it more difficult to activate the thin filaments to allow force production. By targeting this mechanism specifically, using a thin-filament interacting peptide of myosin binding protein C, we were thus able to rescue MG-induced calcium desensitization.

Our previously published mass spectrometry data provides a possible mechanism for how MG affects tropomyosin movement. We found MG glycation of actin K291 [4], which is within a critical region of actin termed the “A-triad”, a cluster which stabilizes the A-state structure, which is similar to the blocked state [46]. Mutations and modifications to residues in this region have been shown to alter function by either stabilizing or de-stabilizing the A-state [47]. We propose that MG forms an irreversible glycation modification on K291 residue in the A-triad, changing the allosteric interactions between actin and tropomyosin to stabilize the blocked state of tropomyosin and therefore reduce *K*_B_ and myofilament calcium sensitivity (**Supplemental Figure 5**).

The computer model also predicted MG glycation inhibits the crossbridge duty cycle, although the model is unable to differentiate between either a decrease in *f* (crossbridge attachment rate) or an increase in *g* (crossbridge detachment rate). The stopped flow and tension cost data both indicated that the detachment rate was unchanged. Combined with the model, these data highly suggest that MG decreases the crossbridge attachment rate. Unfortunately, measuring the attachment rate requires proteins in saturated conditions (i.e., non-physiological), and is challenging to measure with rigorous approaches, so this cannot be confirmed at present. However, one possible explanation for how MG could decrease the attachment rate is by affecting the super-relaxed state of myosin [48]. If MG enhances the super-relaxed state and fewer myosin heads are available to form crossbridges, this could appear as a reduced rate of attachment (*f*). The stopped-flow experiments do not include the impact of the super-relaxed state, so it is not possible to account for its effect. Furthermore, while the tension cost experiments presumably do include the super-relaxed state, myosin heads in this state have a very low basal ATPase activity [49] and might have little effect on the measured tension cost. Thus, whether MG alters the super-relaxed state remains an unaccounted-for possibility in this study.

Our results in myocytes not exposed to MG agree with prior studies showing OM increases myofilament calcium sensitivity [36] and reduces F_max_ likely by depressing the myosin working stroke [50]. We showed that MG blocked the calcium sensitizing effect of OM but did not affect its ability to decrease F_max_, indicating that MG causes OM to act as a negative inotrope. Our finding that MG keeps tropomyosin in the “blocked” position can mechanistically explain this effect. First, OM primes myosin heads to increase their binding to actin to form force-generating crossbridges at submaximal calcium [51], but since MG inhibits thin filament activation, this calcium sensitizing effect is blocked. Furthermore, OM decreases F_max_ by decreasing the myosin working stroke, which would be unaffected by tropomyosin positioning.

Finally, the C0C2 domains of cMyBPC were able to restore the MG-induced decrease in calcium sensitivity. The mechanism of C0C2’s benefit is likely by counteracting the MG-induced reduction in tropomyosin’s blocked-to-closed transition rate, as it has been suggested that C0C2 can alter tropomyosin’s position on actin [18, 52]. These findings provide a strong proof-of-principle that specifically targeting the molecular mechanism of MG can rescue function. However, the C0C2 peptide was not able to restore the decrease in F_max_ caused by MG. The inability of C0C2 to restore F_max_ supports the model prediction that there are two separate effects of MG glycation on the sarcomere, and C0C2 only affects one of these. Whether this second effect is a decrease in the crossbridge attachment rate as the model and experiments suggest and if this dysfunction can likewise be reversed will have to be determined in future studies. Furthermore, whether the C0C2 cMyBPC fragment could be an appropriate therapeutic option in the clinic is not clear. The C0C2 peptide contains the regulatory region of cMyBPC that includes phosphorylation sites that are important for the sarcomere’s response adrenergic stimulation [43, 53, 54], so the peptide would be susceptible to modulation by phosphorylation *in vivo*. However, a recent study expressed the C0C2 peptide using AAV, and showed it was capable of rescuing function in a cMyBPC knockout mouse [55], suggesting the peptide is stable and functional *in vivo*.

## Conclusion

Overall, this study provides a proof of principal that either C0C2 or another therapeutic that can move tropomyosin towards the “open” state would be a potential therapeutic avenue for patients who have diabetes and either have heart failure or are at risk of developing it. As initial treatment approaches failed, such as AGE-breakers, it was critical that we identified this specific mechanism of MG glycation using biophysical, computational, and chemical kinetics. Furthermore, as no small molecules currently target tropomyosin, these results provide rationale that future drug discovery efforts should explore this mechanism. These insights provide a strong foundation for future work targeting these mechanisms to stem the increase in heart failure cases that will occur as the prevalence of diabetes continues to grow worldwide.

## Abbreviations

AG: Aminoguanidine
AGE: Advanced Glycation Endproducts
cMyBPC: cardiac myosin binding protein C
*f*: myosin binding rate
F_max_: maximal calcium-activated force
*g_xb_*: myosin detachment rate
HF: Heart Failure
k: ATP detachment rate constant
*K*_B_: blocked-to-closed rate transition
*k*_tr_: rate of force redevelopment
LV: Left Ventricle
MG: Methylglyoxal
OM: Omecamtiv Mecarbil

## Author contribution

M.P. and J.A.K. designed the project and research studies. M.P., T.K., S.K.B. and S.G.C. conducted experiments and acquired data. M.P., S.K.B., S.C.G., M.J.G. and J.A.K. interpreted experimental results. M.P., S.G.C. and J.A.K. analysed data and created figures. P.P.R., D.L. and S.P.H. provided peptide and human samples. M.P. and J.A.K. wrote the manuscript. T.K., S.K.B., S.G.C., D.L., P.P.R., S.P.H. and M.J.G. revised the manuscript.

